# Rapid spread of African Swine Fever across Borneo

**DOI:** 10.1101/2024.06.20.597708

**Authors:** Olivia Z Daniel, Sui Peng Heon, Christl A Donnelly, Henry Bernard, C. David L Orme, Robert M Ewers

## Abstract

African Swine Fever (ASF) reached the island of Borneo at the end of 2020. The first mortalities occurred in wild bearded pigs (*Sus barbatus*) in Sabah, north-east Borneo. The virus then began to spread across the island but, confounded by COVID 19 lockdowns the spread was difficult to monitor on the ground. The Babi Hutan Project was launched in April 2021 to gather data on pig sightings using citizen science. Here we bring together the data from this project and other online sources to show how the virus encompassed almost the entire island within a one-year period. The speed of spread appears to increase with time following an exponential model: we estimate an average speed of spread of 0.89km/day after 100 days since the first observation and at 4.28km/day after 400 days. We include recommendations of next steps for the bearded pigs of Borneo.

---

African Swine Fever (ASF) is a highly contagious viral haemorrhagic disease affecting members of the family Suidae, (pigs, hogs and boars). In its acute form it can cause 95-100 % mortality ^1–3^, there is currently no approved vaccination and no cure ^3,4^. It is a notifiable disease for the World Organisation for Animal Health (OIE), and documentation of its spread is regularly updated on the Food and Agriculture Organisation of the United Nations (FAO) website ^2,5,6^. Fortunately, it is not zoonotic.

Prior to the ASF outbreak in Borneo, bearded pigs (*Sus barbatus*) were listed as Vulnerable by the IUCN Red List of endangered species ^7^, and are recognised as ecologically and economically vital^8^. Bearded pigs are cited as ‘ecosystem engineers’ dispersing and predating tree seeds, rooting through soil and removing saplings ^9,10^. On Borneo, traditional hunting of bearded pigs dates back over 35,000 years^11^, and before this ASF outbreak their meat represented 50 to 75 % of hunted animal biomass, comprising an important protein source for many non-Muslim communities^8,12,13^.

In Sabah, an unusually high number of dead bearded pigs were observed from December 2020 in Kinabatangan^14^. ASF was confirmed to be the cause of pig mortalities by OIE in February 2021, in Pitas ^6,15^. Reports of bearded pig carcases continued until July 2021 (Fig. 1a). Commercial pig farms confirmed cases in February 2021 with outbreaks reported until October 2022 (Fig. 2a). ASF was first reported in Kalimantan (Indonesian Borneo) in May 2021 ^16,17^ and Sarawak (Malaysian Borneo) in July 2021 ^18,19^ (Fig 1a.). No cases have officially been reported from Brunei.

**Figure 1.**
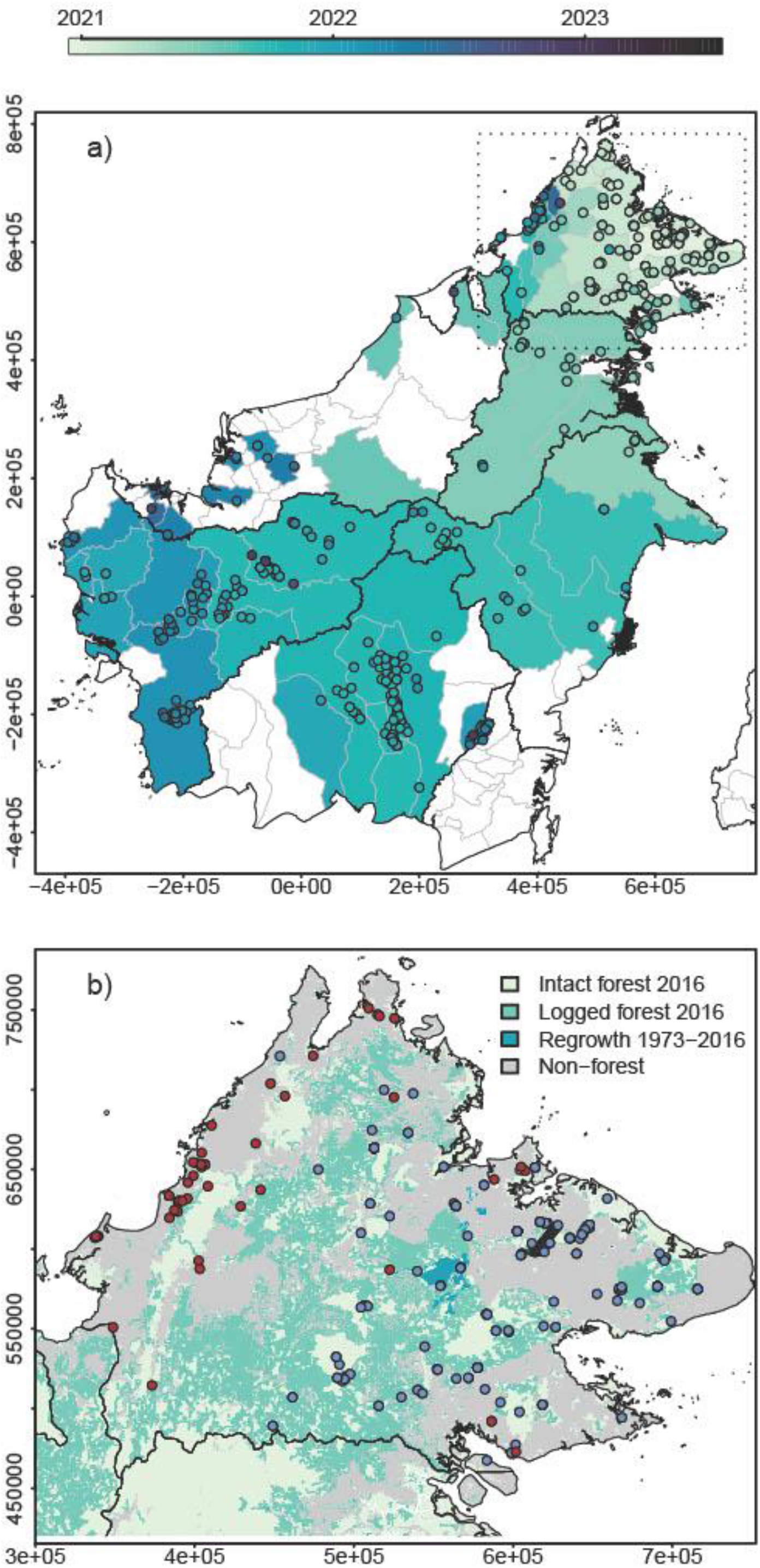
(a) Map showing the first occurrences of ASF in districts in Borneo, colour co-ordinated according to month of detection. All reports of pig deaths due to ASF that had exact locations are superimposed. The lightest shading is for the earliest reports in December 2020 getting darker as time goes by with the last report in July 2023. This map incudes data from both domestic and bearded pigs. (b) Detailed view of forest cover ^21^ and instances of bearded pig (blue points) and domestic pig (red points) mortality in Sabah between Dec 2020 and October 2022. Administrative boundaries are from GADM 4.1.

**Figure 2.**
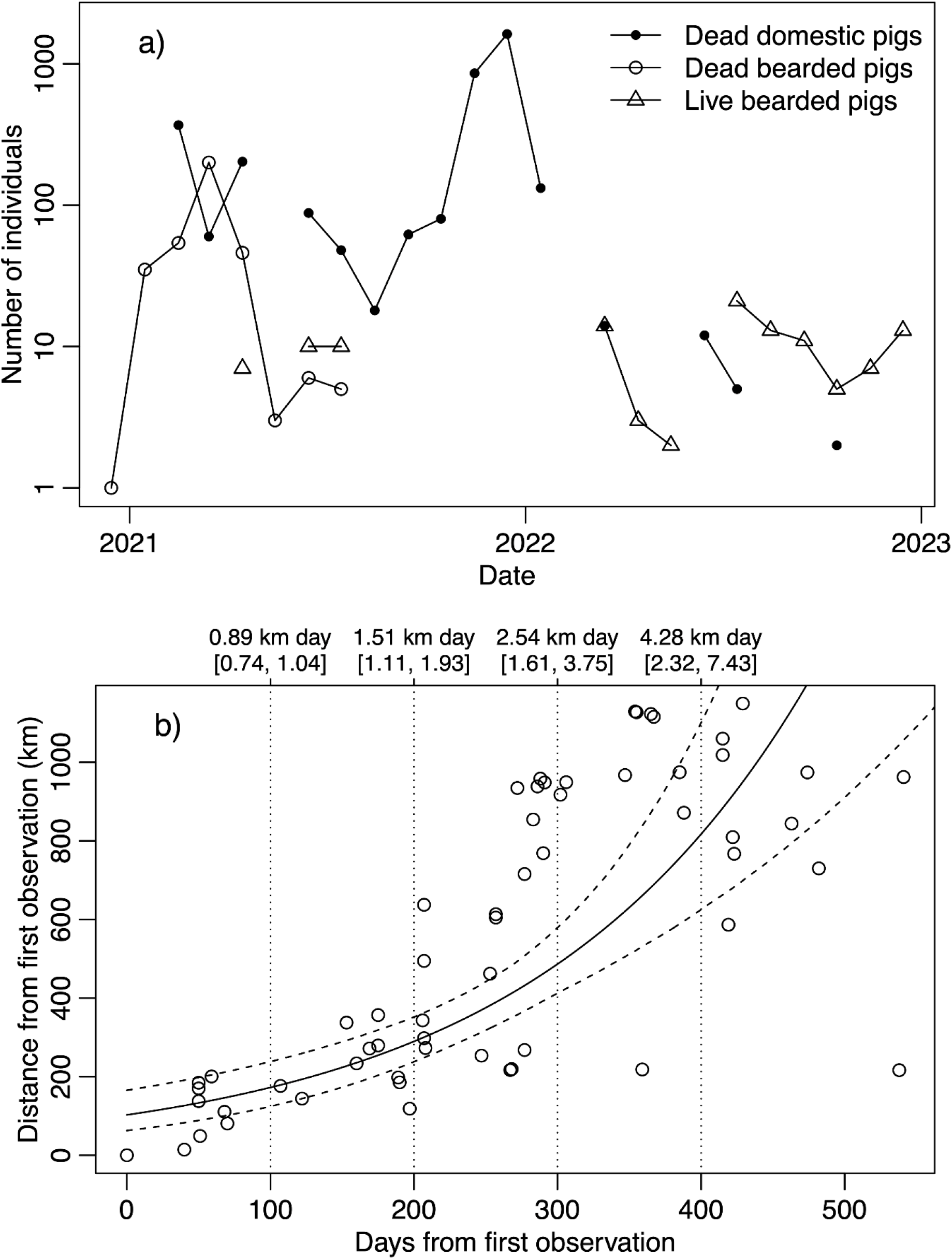
(a) Reports of numbers of live and dead bearded pigs and dead domestic pigs in Sabah, Malaysian Borneo Between Dec 2020 and Dec 2023. Data collated by the Babi Hutan Project ^14^. (b) Scatterplot of date of first reported death by district and distance from the first observation (the date of the first reported death in each district is either the first observation or the earliest reported date at a district level, using mid-month if only the month was recorded). The solid fitted line shows the linear model and 95% CI (dashed lines). The vertical dashed lines are used to indicate the speed of spread (slope of the fitted model) and 95% confidence limits for four time points.

To date, the OIE has only published data on ASF cases in Sabah and Sarawak, and the majority of these document domestic pigs ^6^. Reports of pig deaths in Indonesian Borneo are listed on the FAO website, but also refer almost entirely to domestic pigs ^2^. In addition, iSIKHNAS is an animal health information system for Indonesia, which has reports of ASF cases, although it does not specify whether they are bearded or domestic pigs^20^. Historically, bearded pig populations on Borneo have not been directly monitored, further complicating collating data on the impact of ASF. Limited access to field sites as a result of Covid-19 restrictions also precluded formal rapid response surveillance. Therefore, in April 2021 the Babi Hutan Project was launched (www.babihutan.com), a collaboration between Sabah Wildlife Department, Sabah Veterinary Service, and Imperial College London to track the spread of ASF through wild bearded pig populations using citizen science ^14^. Sightings of bearded pigs, either dead or alive, were requested via the website, social media, and a WhatsApp hotline. These data, along with OIE, FAO and iSIKHNAS reports, have been compiled into a single database which includes both bearded and domestic pig deaths. To date we have collated just over 660 reports of pig deaths across Borneo, amounting to hundreds of bearded pigs and over 150,000 domestic pigs.

Our data indicates that ASF swept rapidly through Borneo expanding from north-east Sabah to the southern districts in Central Kalimantan, and districts on the south-west coast of West Kalimantan, in a period of just 12 months from when the virus was officially recognised (Fig. 1a), although it is possible the spread was quicker. Although we cannot discount the possibility that this rapid spread was aided by multiple incursions of ASF, Khoo et al., (2021) demonstrated that the ASF virus found in three locations in Sabah were of the same strain which matches that in the Asia-Pacific region and indicates a common origin^22^. Within Sabah, the first wave of ASF affected bearded pigs from December 2020 to June 2021 and the second wave from October 2021 to January 2022 affecting domestic pigs (Fig. 2a). Between February 2021 and January 2022, 72 outbreaks of ASF were reported to OIE in Sabah, and no further cases of ASF have been reported to OIE in Sabah since July 2022 ^6^. In Sarawak, Malaysian Borneo cases were reported in areas bordering Sabah and North Kalimantan in July 2021 with a second wave of cases in January 2022 with the latest outbreaks reported in October 2022 (Fig 1a.). Cases began to appear in Indonesian Borneo in May 2021 in both North and East Kalimantan and in September 2021 in both Central and West Kalimantan, the most recent outbreak was in July 2023^17^ (Fig 1a). There have been no cases reported in South Kalimantan.

Assuming ASF on Borneo did all originate from the first observation in the north-east of Sabah, it is possible to model the rate of spread from that date and that location (Fig. 2b). We used a linear model to fit log distance from first observation (d) as a function of the time since the first observation (t), generating an exponential model of the rate of spread. The model has good explanatory power (adjusted R2 = 0.54) and the slope and intercept are both highly significant (p << 0.0001). The modelled equation parameters and standard errors are `d = exp(4.63 [±0.186] + 0.00519 [±0.000631] t)`. The slope of this equation gives the speed of spread at a given time, and we refitted the model to bootstrapped data (n=100000) to estimate the 95% confidence limits around both the fitted model and the estimated speed oof spread for four time points during the epidemic (Fig. 2b).

To date, there are no formal estimates quantifying the magnitude of the population crash that beaded pigs have suffered due to ASF. Officially reported cases only represent ASF cases verified by positive laboratory test results. It has been estimated that 90 % of Sabah’s bearded pig population has disappeared ^23^, which is in line with ASF outbreaks in other wild pig populations ^24^. There are anecdotal reports that the situation may be similar in Sarawak’s bearded pig populations ^25^. Reports of pig footprints and wallows are starting to come in as well as sightings of low numbers of live bearded pigs in Sabah (Fig. 2a), these include sightings of juveniles, indicating that survivors have reproduced.

Hunting licences for bearded pigs have been suspended in Sabah since February 2021 ^26^. It is essential that the current ban is enforced and that educational materials are made available to ensure that those that usually hunt the bearded pigs are aware of the ban that is in place and why. It is also essential to ban the sale of bearded pig meat, and to educate the general public to avoid consumption.

Continued implementation of the already existing ASF prevention protocols^27^ - including controlled movement of pigs and pork products between states and countries, awareness campaigns, and close monitoring activities in farms and abattoirs in all affected districts as well as enhancing biosecurity measures – is also key. For diseases such as ASF which have no cure, prevention of spread is key, therefore any pig death or even pig illness should be investigated immediately^3^. Moreover, particular care should be taken to ensure ASF does not reach the bearded pig populations on the smaller islands that surround Borneo and indeed other islands with endemic pig populations, as Southeast Asia represents an important area for pig diversity with 11 endemic species found in the region^8,28^.

Finally, we suggest a research agenda that can help guide further decision-making regarding control of the ASF outbreak. The continued collection and collation of bearded pig sightings and monitoring data are urgently required to obtain up-to-date information on the abundance and age structure of the remnant population. Such data are needed across the breadth of Bornean land uses, including protected areas, managed agricultural land, and forest remnants in the vicinity of domestic pig farms. We urgently need confirmation, via blood samples, as to whether bearded pigs have survived because they have been resistant or because they have evaded infection with the virus responsible ASF. It would also be prudent to collect any ectoparasites from captured pigs to determine if they contained any trace of the ASF virus and could act as potential vectors. ASF disease transmission models^29^ should be developed and adapted to the Bornean landscape to help identify potential refugia and to guide further decision making processes. Finally, the IUCN Red List status of the bearded pig should be urgently re-evaluated^8^.

We are cautiously optimistic that ASF has not spelt the end for the bearded pigs of Borneo. We hope that if the survivors are given the chance to repopulate, that these engineers of Borneo’s forests can resume their work.

## Acknowledgements

We thank Dr Peter Lee (Sabah Veterinary Service) and Dr Sen Nathan (Sabah Wildlife Department) for their support in establishing the Babi Hutan website. We are very grateful to Department of Veterinary Services, Sabah Wildlife Department, Sabah Forestry Department, Maliau Basin Conservation Area, Danum Valley Field Centre, Danau Girang Field Centre, HUTAN, Wildlife Rescue unit, Yayasan Sabah, BORA (Bringing Back Our Rare Animals), PANTHERA, Rhino and Forest Fund, SAFE Project, 1StopBorneo, Borneo Mammal Club, South East Asia Rainforest Research Partnership, The Orangutan Foundation and the Bornean Sun Bear Conservation Centre for sharing their sightings. Celina Chien, Jeffery Chan, Josh Jones and Vivian Hughes provided technical support.

## Author contributions

OZD wrote the article with assistance from SPH, HB, CAD, RME and CDLO. OZD and SPH were primarily responsible for the collation of the data with supervision from RME. HB and CAD provided guidance on recommendations for future work. CDLO, CAD and OZD undertook data analysis. All authors have given final approval for publication of this work and agree to be accountable.

## Declaration of interest statement

The authors report there are no competing interests to declare.

## Notes

https://zenodo.org/records/10568985

### Competing Interest Statement

The authors have declared no competing interest.

